# Resolving the spatial profile of figure enhancement in human V1 through population receptive field modeling

**DOI:** 10.1101/2019.12.18.881722

**Authors:** Sonia Poltoratski, Frank Tong

**Author notes:** Correspondence addressed to: Sonia Poltoratski *Current address:* 450 Serra Mall, Building 420 / Jordan Hall, Stanford, CA 94305, *Email:*.

## Abstract

The detection and segmentation of meaningful figures from their background is a core function of vision. While work in non-human primates has implicated early visual mechanisms in this figure-ground modulation, neuroimaging in humans has instead largely ascribed the processing of figures and objects to higher stages of the visual hierarchy. Here, we used high-field fMRI at 7Tesla to measure BOLD responses to task-irrelevant orientation-defined figures in human early visual cortex, and employed a novel population receptive field (pRF) mapping-based approach to resolve the spatial profiles of two constituent mechanisms of figure-ground modulation: a local boundary response, and a further enhancement spanning the full extent of the figure region that is driven by global differences in features. Reconstructing the distinct spatial profiles of these effects reveals that figure enhancement modulates responses in human early visual cortex in a manner consistent with a mechanism of automatic, contextually-driven feedback from higher visual areas.

**Significance Statement:** A core function of the visual system is to parse complex 2D input into meaningful figures. We do so constantly and seamlessly, both by processing information about visible edges and by analyzing large-scale differences between figures and background. While influential neurophysiology work has characterized an intriguing mechanism that enhances V1 responses to perceptual figures, we have a poor understanding of how the early visual system contributes to figure-ground processing in humans. Here, we use advanced computational analysis methods and high-field human fMRI data to resolve the distinct spatial profiles of local edge and global figure enhancement in the early visual system (V1 and LGN); the latter is distinct and consistent a mechanism of automatic, stimulus-driven feedback from higher-level visual areas.

## Introduction

When we view a scene, our visual system must detect and segment figures from the background environment, guiding our attention toward regions that are likely to contain meaningful objects. Research has shown that local differences in visual feature content (e.g., color, luminance, or orientation) between a figure and its surroundings provide powerful cues for segmentation and detection (Treisman and Gelade, 1980; Bergen and Adelson, 1988; Landy and Bergen, 1991; Nothdurft, 1993). Conversely, if the figure shares many visual features with its surrounding background, it may be much more difficult to detect; this property of vision is commonly exploited by animals whose coats or skins match their typical environment, making them more difficult for predators to see (Figure 1A). How might the contextual processing of features at local and larger scales contribute to the visual perception of figures?

**Figure 1.**
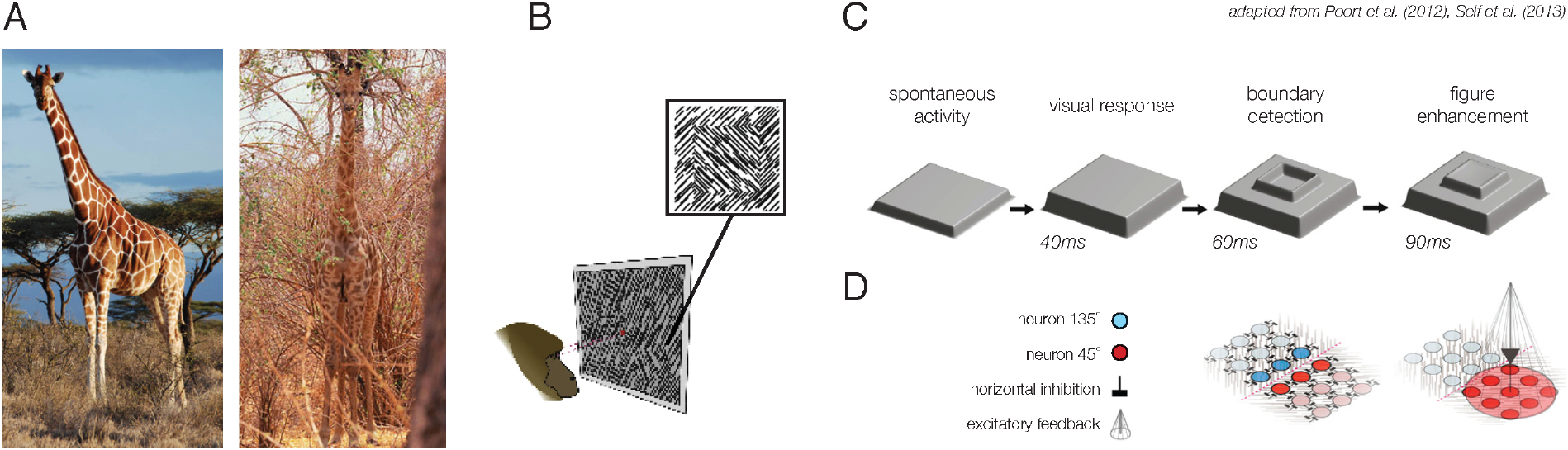
Figure-ground modulation in the wild and in the literature. (A) An example of animal camouflage, which is pervasive in the natural visual world. Detection of figures is much more difficult when they resemble the color, luminance, and spatial frequency content of the surrounding background. (B) A sample orientation-defined figure display, typically shown to an animal such that the recorded neuron’s receptive field falls over the figure region. (C). Proposed mechanisms of figure-ground modulation in the primate V1: after an initial visual response (40ms), local boundary detection mechanisms enhance responses near the figure-surround boundary (60ms). Subsequently (90ms), the entire figure region is enhanced. (D) The contribution of intrareal horizontal inhibition and excitatory feedback from higher-level visual areas to boundary detection (left) and figure enhancement (right). *(B-D) adapted from Poort et al. (2012) and Self et al. (2013); to request permission upon manuscript acceptance*.

Researchers have sought to determine the early visual mechanisms by which figures are differentiated from a surrounding background (i.e. figure-ground perception), with considerable focus on the response properties of neurons in the primary visual cortex (V1) of non-human primates (Lamme, 1995; Zipser et al., 1996; Super et al., 2001; Marcus and van Essen, 2002; Self et al., 2013; 2019). These studies find evidence of two distinct mechanisms by which figures are differentiated from ground regions in primary visual cortex (V1): when a feature-defined figure, such as a patch of oriented lines, is presented together with a background region (Figure 1B), V1 neurons with receptive fields that fall along the boundary between figure and ground exhibit a rapid early enhancement in their response (Figure 1C, “boundary detection”). Subsequently, V1 neurons corresponding to the figure region itself exhibit response enhancement during the sustained phase of the neural response (Figure 1C, “figure enhancement”). The finding of this delayed figure enhancement in primary visual cortex is of particular interest, as it suggests the involvement of more complex neural processes that cannot readily be explained in terms of local feature-tuned surround suppression (Sillito et al., 1995; Bair et al., 2003; Shushruth et al., 2012; Bijanzadeh et al., 2018), particularly by its asymmetrical spatial profile. Indeed, subsequent electrophysiology work has suggested that top-down feedback from higher-order visual areas, including area V4 and the parietal cortex, is likely responsible for inducing figure enhancement responses in V1 (Lamme et al., 1998a; Poort et al., 2012; Self et al., 2013; Poort et al., 2016).

Recently, our group has found that feature-defined perceptual figures evoke enhanced responses throughout the human early visual system, including V1 and the lateral geniculate nucleus (LGN) of the thalamus (Poltoratski et al., 2019). While figure-ground modulation has been well-characterized in the early visual cortex of non-human primates, in humans the processing of textures, surfaces, and shapes has often been attributed to cortical areas beyond the primary visual cortex, including V4 (Kastner et al., 2000; Kourtzi et al., 2003; Thielscher et al., 2008) and the lateral occipital cortex (Grill-Spector et al., 1999; Vinberg and Grill-Spector, 2008). Our recent work demonstrated the involvement of human primary visual cortex in the enhancement of visual figures (Poltoratski et al., 2019); importantly, we showed that figure-ground modulation in human early visual cortex and LGN is dissociable from effects of directed attention, suggesting an automatic, contextually-driven process. However, neurophysiological studies in monkeys have suggested that figure enhancement in V1 becomes severely attenuated if the animal must attend elsewhere to perform a visual task, while border responses remain intact (Schira et al., 2004; Poort et al., 2012). Could it be that our previous observations of enhanced responses to figures in human V1 was caused by local responses specific to the border between the figure and the surround, or can we find evidence of a separate, spatially specific enhancement of the figure region?

One can differentiate between responses driven by the processing of local differences between figure and surround from those caused by a more global figure enhancement by measuring their spatial profiles, which are predicted to be markedly different: boundary responses should appear relatively local to the border between figure and surround whereas figure enhancement should spread across the entire figure region (e.g., Zipser et al., 1996; Self et al., 2019). The prevailing theory of figure enhancement predicts that retinotopic enhancement of activity should be evident throughout the extent of the figure, and moreover, this enhancement should not vary as a function of distance from the boundary (c.f. Poort et al., 2012). However, others have challenged the idea that the full extent of the figure may be enhanced in V1, especially at locations far from the boundary (Rossi et al., 2001; Zhaoping, 2003).

## Materials and Methods

The goal of our study was to determine whether figure enhancement does indeed occur in human V1 in a manner that can be distinguished from boundary responses, and to determine how the magnitude of this enhancement changes across the figure region as a function of distance from the boundary. We relied on population receptive field (pRF) mapping (Dumoulin and Wandell, 2008; Kok and de Lange, 2014; Wandell and Winawer, 2015) to characterize the response fields of individual voxels in order to distinguish figure enhancement from local responses elicited by the boundary between figure and surround. Observers performed a visual task (color change detection) at central fixation while task-irrelevant figure-ground stimuli (4° or 6° in diameter) were presented (Figure 2A, Movie 1). We compared the spatial profile of fMRI responses to 3 types of visual displays: a *ground-only* condition in which no figure was present, a *congruent figure* condition in which the figure region is iso-oriented with the background, resulting in a weak figure percept, and an *incongruent figure* condition in which the figure is orthogonal to the background and appears more perceptually distinct.

**Figure 2.**
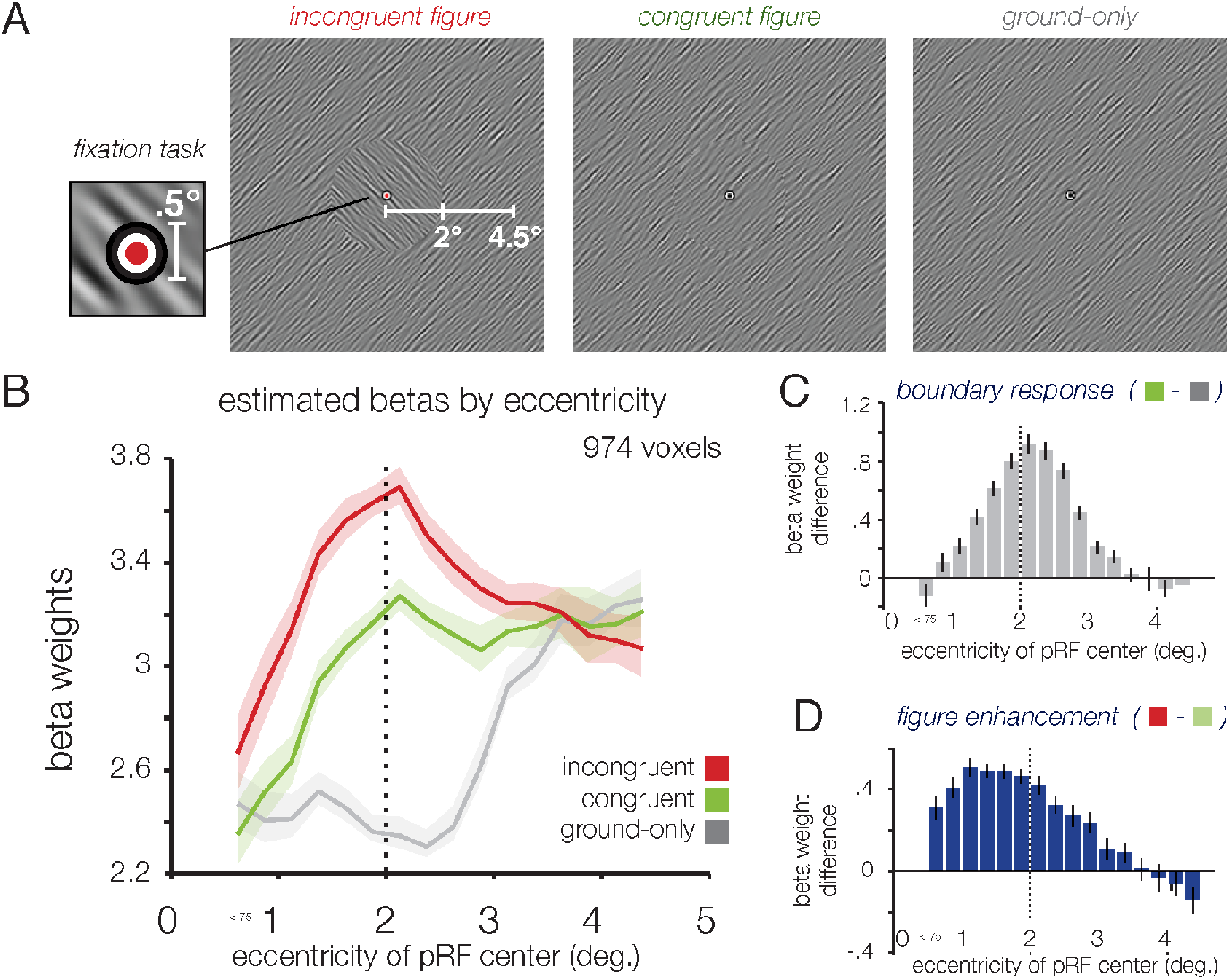
Boundary responses and figure enhancement as a function of eccentricity. (A) Examples of spatially filtered oriented-noise stimulus displays used in the Experiment 2. Three main conditions are depicted: (left) incongruent figure, in which the figure and surround were orthogonally oriented, creating both a perceptual border and an orientation difference; (middle) congruent figure, in which the figure and surround are iso-oriented but sampled from distinct patches of noise, creating a visible phase-defined border but a weaker figure percept; (right) ground-only, in which a single oriented texture fills the visual field. The spatial frequency depicted here is much lower than the experimental stimuli (0.5-8 cycles per degree) to improve visibility. See also Movie 1. (B) V1 responses plotted using mean Beta weights for each experimental condition as a function of the eccentricity of each voxel’s pRF center. Dotted line at 2° eccentricity indicates the figure-surround border. (C) Top: V1 BOLD Beta weight difference between congruent figure and ground-only conditions, associated with a predicted boundary response. Bottom: V1 BOLD Beta weight difference between incongruent and congruent figure conditions, associated with a predicted figure enhancement due to differences in orientation. The spatial profile of this effect as a function of eccentricity is clearly distinct from that of the top panel, and appears to extend through the full eccentricity range of the figure but to decline beyond the 2° figure-surround border.

While we expected V1 responses to be greater overall for both types of figures than when no figure is presented (ground-only condition), we were particularly interested in how responses to the incongruent figure would differ from the congruent condition in their spatial profile. We recently found that V1 responds more strongly to incongruent figures than to congruent figures, even when attention is directed away from the perceptual figure (Poltoratski et al., 2019; see also Marcus and van Essen, 2002). However, to conclude that this modulation is driven by figure enhancement, it is necessary to resolve the spatial profile of local, spatially constrained boundary responses, which we predicted would occur in response to the congruent figure stimulus, and to demonstrate that an additional enhancement occurs throughout the greater figure region, as we predicted would occur for incongruent figures. Through pRF mapping and the visual reconstruction of fMRI responses to incongruent figures as compared to congruent ones, we find compelling evidence that figure enhancement occurs in human V1 in a manner that can be clearly distinguished from local boundary processing.

### Participants

All experiments were performed at the Vanderbilt University Institute for Imaging Science (VUIIS), adhering to the guidelines of the Vanderbilt IRB; participants were compensated for their time. Six experienced human observers participants (four females) ages 23-31 were scanned in the main experiment; three of these participants (all females) also completed an additional control experiment, in which larger figures (6° in diameter) were presented. Author SP participated in all experiments.

### Scanning procedure

All functional data were collected at the VUIIS research-dedicated 7 Tesla Philips Achieva scanner using a quadrature transmit coil in combination with a 32-channel parallel receive coil array. BOLD activity was measured using single-shot, gradient-echo echoplanar T2*-weighted imaging, at 2-mm isotropic voxel resolution (40 slices, TR 2000ms, TE 35ms; flip angle 63°; FOV 224 x 224; SENSE acceleration factor of 2.9; phase-encoding in anterior-posterior direction). Each MRI session lasted 2 hours, during which we acquired the following images: i) 1-2 functional localizer runs using a central flickering checkerboard to identify retinotopic regions in visual cortex and LGN that corresponded to the stimulus location ii) 7-8 fMRI runs to measure BOLD activity during the experiment, and iii) 5-8 fMRI runs to map population receptive fields in these voxels. Each of the run types lasted 4-6 minutes.

### Scanning procedure: Retinotopy

Prior to participating in the functional experiments, each participant underwent a separate session of retinotopic mapping. We used a typical phase-encoded design (Engel et al., 1997; Wandell et al., 2007) in which subjects fixated while they viewed flickering checkerboards consisting of rotating wedges to map polar angle and expanding rings to map eccentricity (Swisher et al., 2012). Retinotopy data were acquired at the VUIIS using a Philips 3T Intera Achieva MRI scanner equipped with an 8-channel receive coil array, using 3-mm isotropic resolution (TR 2s, TE 35ms, flip angle 80°, 28 slices, 192 x 192 FOV).

### fMRI preprocessing

Data were preprocessed using FSL and Freesurfer tools (documented and freely available for download at http://surfer.nmr.mgh.harvard.edu), beginning with 3D motion correction and linear trend removal, followed by slice-timing correction for pRF runs and a high-pass filter cutoff of 60s. Functional images were registered to a reconstructed anatomical space for each subject; this registration was first automated in FSL and then checked and corrected by hand. This allowed for the alignment of the fMRI data to the retinotopy data, which was collected in a separate session. The functional localizer data was spatially smoothed using a 1-mm Gaussian kernel to improve the spatial contiguity of delineated regions of interest; no spatial smoothing was performed on the experimental or pRF mapping runs. Further analyses were conducted using a custom Matlab processing stream. Each voxel’s intensities were normalized by the mean of the time series, converting to mean percent signal change within each run. Outliers were defined as time points for which the voxel’s response measured more than 3 times its standard deviation from its mean, and were Winsorised (Hastings et al., 1947). This condition-blind preprocessing step minimizes the impact of rare spikes in MR intensity while preserving the temporal structure of the responses in each voxel.

In all functional experiments in this study, we relied on an fMRI block paradigm, and presented blocks of visual stimulation (16s duration) interleaved among 16s fixation-rest periods. The amplitude of the BOLD response during each stimulus block was then estimated using the general linear model for each voxel. These estimates were averaged across blocks by experimental condition to yield voxel-wise mean standardized Betas in each condition.

### ROI localization and voxel selection

To initially define retinotopic visual areas V1-V3, each subject participated in a separate retinotopic mapping scan. Boundaries between retinotopic areas V1-V3 were delineated by hand, by identifying reversals in the phase of the polar angle map measurements. The resulting labels were aligned to the functional space of the current experiment using FSL and Freesurfer software, and this registration was checked and corrected by hand. Additionally, 1-2 runs of functional localizer, described below, were collected in the main experimental sessions to identify the LGN in each subject and to select regions of interest (ROIs) responsive to our experimental display from the V1-V3 retinotopic areas. The localizer consisted of blocks of a flickering checkerboard stimulus spanning the full 9° field of view, and was designed to yield a large ROI that could be then refined by pRF model fitting.

For all analyses, we used these functional labels in conjunction with the pRF fitting results to define regions of interest. For each subject, all voxels in each visual area were fitted with the pRF model as described below. For further analyses, we used voxels whose pRF centers were within the range of the mapping stimulus (0.25° to 4.5° eccentricity) and were larger than 0.1°; this limit trimmed instances in which the model predicted nearly no visual response to the mapping stimulus. Following this trimming procedure, we selected the top 33% of best-fitted voxels for each subject in each ROI, as indexed by the R^2^ between observed and predicted data. In V1, this yielded fits with R^2^ cutoffs that ranged from 0.62 to 0.81 in individual subjects (mean = 0.71); corresponding V1 ROIs for each subject ranged from 131 to 187 voxels bilaterally (mean = 162). In the LGN, R^2^ cutoffs ranged from 0.16 to 0.27 (mean = 0.22), yielding ROIs that were 17-36 voxels in size (mean = 26.5).

### Population receptive field mapping

Population receptive fields (pRFs) correspond to the location in visual space that best drives activity in the population of neurons in each voxel (Dumoulin and Wandell, 2008; Wandell and Winawer, 2015). Unlike standard retinotopic mapping, pRF mapping resolves not only the central location that best drives responses in each voxel, but also the spatial extent of this response field. In each fMRI experimental session, we mapped population receptive fields in retinotopic areas V1-V3, using a 2D circular Gaussian model of pRF structure. PRF modeling involves presenting a mapping stimulus that spans the visual field over time, and estimating the pRF parameters that most likely produced the measured BOLD response in a particular voxel. These parameters define the pRF’s location, size, and gain. pRF properties for each voxel are assumed to reflect the combined RFs of the neural population in a voxel, and appear well-aligned with single-neuron receptive field properties, such as contralateral preference, increasing size at greater eccentricities, and increasing size as one ascends the visual hierarchy (Dumoulin and Wandell, 2008; Wandell and Winawer, 2015). Here, we mapped pRFs using a traveling-bar stimulus comprised of rapidly-presented, full-color objects embedded in pink noise (developed by Kendrick Kay, *kendrickkay.net/analyzePRF*; object stimuli from Kriegeskorte et al., 2008). The bar stimulus swept through a circular region with 4.5° radius, the maximal visible field of view at our 7T scanner, and each mapping run lasted 5 minutes. Voxel-wise responses for each visual area were fitted with a 2D Gaussian pRF model using a custom Matlab pipeline. We used a dual-stage multidimensional nonlinear minimization (Nelder-Mead) fitting procedure: each voxel was initially fitted in a downsampled stimulus space with a fixed Gaussian σ (1°), and then these parameters were used to initialize a full model fitting in native stimulus space. Estimated parameters described each voxel’s pRF position (X, Y), size (σ), and response amplitude; we can additionally convert these parameters to measures of polar angle, eccentricity, or full-width half-maximum (FWHM) to convey pRF size.

### Reconstruction of spatial profiles of figure-ground modulation

For the spatial profile visualization, pRF mapping was used to project the differential responses evoked the figure-ground stimuli to stimulus space. Previous studies (e.g. Kok and de Lange, 2014) have adopted an approach of projecting the weighted activity of each voxel into image space by scaling its Gaussian pRF by the voxel’s response to the display, and then calculating the linear sum of all pRFs in a region. In our work, we have found that a multivariate regression-based approach leads to sharper projections. Specifically, we used ridge regression (Hoerl and Kennard, 1970) as described in Figure 7: we model the voxel-wise responses in each region as a linear product of those voxels’ pRFs and the spatial profile of perceptual effects driving these responses. The spatial profile is thus solved as a matrix of predictors (*β*) in the regression, with each weight representing a portion of the original stimulus space (downsampled to 0.25° per weight/pixel, or 4 pixels per degree; the pRFs are similarly subsampled). The resulting regression can be described as a standard linear model: *y* = *Xβ* + *ε*, where *y* ∈ ℝ^*N*^ represents the vector of voxel responses, *X* ∈ ℝ^*N*×*P*^ consists of a 2D matrix of each voxel’s mapped Gaussian pRF, *β* ∈ ℝ^*p*^ are the predictors (pixels) to be estimated, *ε* is the error term, and *p* and *N* are pixels and voxels, respectively. Since there are many more predictors (pixels) than voxel responses to be predicted, this leads to an inverse mapping problem, for which standard multivariate regression cannot find a unique solution. To constrain the estimation procedure and to minimize overfitting, regularized ridge regression can be used to estimate the predictors for each pixel’s contrast. Whereas linear regression seeks to minimize the sum of squared residuals, ridge regression, or L2 regularization, applies an additional penalty term based on the sum of squared weights (*β*) to be estimated, thereby giving preference to solutions with smaller *β* values (Hoerl and Kennard, 1970), as follows: 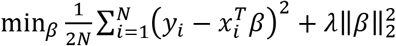, where *λ* is the scaling factor of the penalty term. Here, we performed ridge regression over the data of each individual ROI and subject to produce projection images. The *λ* penalty term for each regression was chosen from the range 0 to 5000 (in increments of 25), using ten-fold cross-validation to find lowest RMSE; in this implementation, a *λ* of zero corresponds to non-regularized regression. The resulting images were z-scored within subjects, averaged, and smoothed with a small Gaussian kernel (σ = 0.75°) to create the spatial profile images in Figure 7.

### Code accessibility

Following the publication of this paper, code for pRF fitting and regression-based visualization will be made publically available at github.com/soniapolt/pRF-figGround.

### Experimental displays

In all experiments, observers viewed orientation-defined figures. To minimize the prevalence of high spatial frequency energy artifacts between the figures and surround, we used oriented bandpass-filtered noise (100% contrast, bandpass filtered from 0.5-8 cycles per degree with a center orientation of 45° or 135°, ~20° FWHM in the orientation domain). Each orientation stimulus was dynamically generated from random white noise, bandpass filtered in the Fourier domain, convolved with a small Gaussian kernel (in the Fourier domain) to minimize Gibbs ringing artifacts, and then converted back to the image domain. The experimental conditions were presented in 16s stimulus blocks, during which the oriented noise patterns were dynamically regenerated every 200ms; these stimulus blocks were interspersed with 16s fixation-rest blocks. A movie of the experimental conditions for the main and control experiments is available (Movie 1).

### Experimental design: Main experiment

Sample displays for each of the three conditions in this experiment are illustrated in Figure 2A. In both the incongruent figure and congruent figure conditions, observers viewed a single, centrally presented figure (4° in diameter). For the congruent condition, figure and surround images were generated from two different noise patterns to create a phase-misaligned figure-surround display. In the main experiment, the center and surround directly abutted (but see Control Experiment A).

In this experiment, participants passively viewed the figure-surround displays while they performed a color-change detection task at fixation throughout the entire experimental run. The fixation changed from black to red for increments of 200ms at random time intervals, occurring on average of 4 times per 16s experimental or fixation block. Participants reported these events by pressing a key on an MR-compatible button box; average percent correct across participants was 94.5% (SD = 5.8%). The task difficulty was titrated for this experiment to avoid load-based suppression around fixation, which we have observed in prior studies (Cohen and Tong, 2015) when more difficult central tasks are used. The figures were not task-relevant at any point in the main experiment nor in the controls; accordingly, we saw no difference in task performance relative to the onset/offset of stimulus blocks (Figure 3), nor were subjects impaired at the task during any of the stimulus conditions relative to blank periods (*F*(3,20) = 0.46, p = 0.72). Overall, we see no positive evidence that participants’ attention was drawn away from the task by the figures, neither generally (Figure 3A) nor in a pattern that would predict the observed enhancement of orientation-defined figures (Figure 3B).

**Figure 3.**
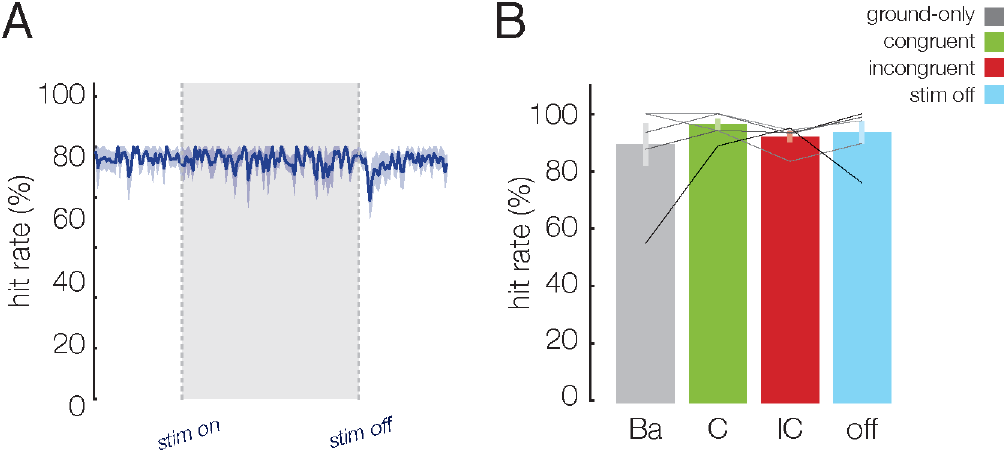
Behavioral performance in the main experiment. (A) Behavioral hit rate relative to the onset and offset of 16s stimulus blocks, plotted in 200ms increments. Performance is averaged across blocks; shaded error bars indicate ± SEM across subjects. There is no apparent difference between task performance when the stimulus is on the screen versus off, save for a slight dip immediately after the stimulus block that is likely attributable to increased blinking. (B) Average hit rate across for targets coinciding with the three main experimental condition stimulus blocks, and stimulus-off blocks. Error bars indicate ± SEM across subjects, and greyscale lines correspond to individual participants’ performance (some overlap). There is no significant effect of stimulus condition on performance of the fixation task (*F*(3,20) = 0.46, p = 0.72), nor is there an overall difference between stimulus-on and stimulus-off performance (*t*(5) = 0.59, p = .58). While the fixation task in these experiments is not particularly challenging, we would expect some variation in performance if the task-irrelevant figures were actively or preferentially attracting subjects’ attention.

### Control Experiment A: gap inserted between figure and ground

In the same session (N=6) as the main experiment, participants also viewed oriented figures that were encircled with a 0.5° greyscale gap (2.0-2.5° eccentricity; illustrated in Figure 8A and Movie 2). This created a visible border around both the incongruent and congruent figures; importantly, the boundaries around these two types of figures were better matched in terms of the local V1 responses they would be expected to evoke as compared to those in the main experiment (which were either phase-defined, or orientation-and-phase-defined). This allowed us to determine whether a difference in local stimulus energy could explain the spatial profile of the effects in the main experiment, and to test whether a visible boundary was sufficient to produce figure-filling when the figure orientation did not differ from the surround. Experimental timing, task, and design were identical to the main experiment.

### Control Experiment B: larger figures

Three subjects from the original experiment performed an additional control experiment in a second session. This control experiment sought to evaluate whether larger figures would still lead to figure enhancement in V1 in regions corresponding to the center of the figure in a manner that could not be readily explained by local effects of feature-tuned surround suppression. This experiment followed the same procedures as the main study, but used figures that were 6° in diameter. Given our limited field of view in the 7T scanner (~9° in diameter), this stimulus precluded us from measuring many voxels that responded primarily to the surround, but allowed us greater measurement of a range of voxels whose pRFs were within the figure.

## Results

### Distinct effects of boundary responses and figure enhancement in V1-V3

Figure 2B shows V1 BOLD responses in each of the experimental conditions binned by the eccentricity of the estimated pRF center for each voxel. Bins are 0.25° wide, and thresholded to contain at least 10 voxels pooled across subjects (see Figure 4 for details of the binning procedure). fMRI responses were generally stronger in the congruent condition than in the ground-only condition, especially around 2° eccentricity, which corresponded to the location of the boundary between the figure and surround. However, fMRI responses were even greater in the incongruent condition, and this differential response could be observed even in voxels with pRFs located near the center of the figure. It should be noted that the spatial extent of our visual display at the 7 Tesla scanner prevented us from examining the responses of voxels whose pRF centers were located much beyond 4° eccentricity; nevertheless, a modest trend of weaker responses in the incongruent figure condition can be seen, consistent with previous reports that perceptual figures can lead to a suppression of neural responses to the ground region (Appelbaum et al., 2006; Poort et al., 2016; Self et al., 2019).

**Figure 4.**
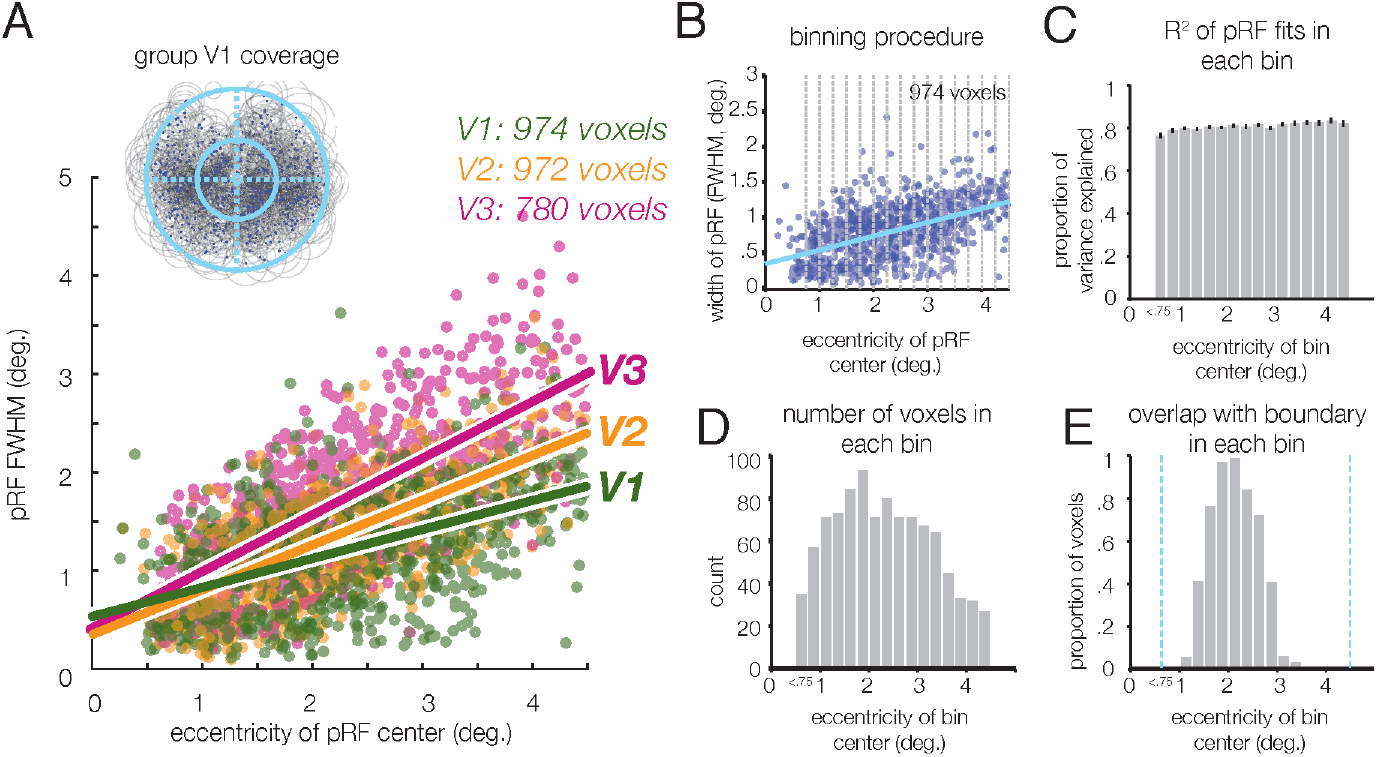
Details of the pRF mapping and binning procedure used in V1 analyses of Figure 2. (A) Estimated pRF size (as measured by Gaussian FWHM) plotted as a function of eccentricity for all voxels used in the analyses for the main experiment. pRF size increases both as a function of eccentricity and as one ascends the visual hierarchy from V1 to V3. Inset: Total pRF coverage of V1 across subjects; blue dots mark pRF centers, and grey outlines mark FWHM. Blue circles indicate the figure location (2° eccentricity) and the full display extent (4.5° eccentricity). (B) Scatterplot of voxel size (Gaussian FWHM) as a function of eccentricity, with grey lines indicating bin edges. Bins were 0.25° wide with the exception of the most foveal, which included all voxels with a center pRF eccentricity < 0.75°. (C) Goodness-of-fit of the pRF model across voxels in each bin. Error bars indicate ± SEM across subjects. (D) Number of voxels in each bin, across subjects. (E) Proportion of voxels in each bin that overlap with the boundary (as defined in Figure 5A), indicating clear differentiation of voxels in the figure, boundary, and surround regions.

We calculated the difference in fMRI response amplitude for congruent figures minus ground-only (Fig. 2C; “boundary response”); this revealed enhanced V1 responses centered at the boundary between the figure and surround as predicted. A modest degree of skewness towards farther eccentricities was also observed, consistent with the fact that pRF sizes generally increase with eccentricity (Figure 4A). This boundary response, or sensitivity to a phase-defined border, appears distinct from the predicted spatial effects of figure enhancement, as it does not spread inward toward the center of the figure.

To determine the spatial profile of the additional enhancement caused by presenting a more distinct figure that differs in orientation from its surround, we calculated the difference in fMRI response for incongruent figures minus congruent figures (“figure enhancement”); this revealed enhanced activity that appeared to extend throughout the full 2°-radius of the figure (Figure 2D). However, as some variability in pRF size is present at each eccentricity (Figure 4A), it is important to consider whether these effects are primarily driven by voxels that, while centered within the figure, nevertheless receive input from the location of the boundary. To address this question, we performed another analysis that took into consideration both the center position and spatial extent of voxel pRFs. Specifically, we evaluated whether the greater response to incongruent figures might be driven by voxels whose pRFs are centered within the figure but also extend over the boundary, in which case local processing of orientation differences might underlie the apparent effect of figure enhancement. We binned the response of voxels according to whether the most sensitive central region of their pRF, defined by the full-width-half-maximum (FHWM) envelope, was contained within the figure (negative values of ‘overlap’ with the boundary) or extended past the boundary (positive values of ‘overlap’), as illustrated in Figure 5. While the predicted boundary response (congruent figures minus ground-only) occurs primarily in voxels whose pRFs overlap with the boundary (gray curve), the magnitude of the figure enhancement effect (incongruent minus congruent conditions, blue curve) remains stable and positive for voxels whose response region (FWHM) falls within the figure and does not overlap with the boundary (F(1,3) = .50, p = .53).

**Figure 5.**
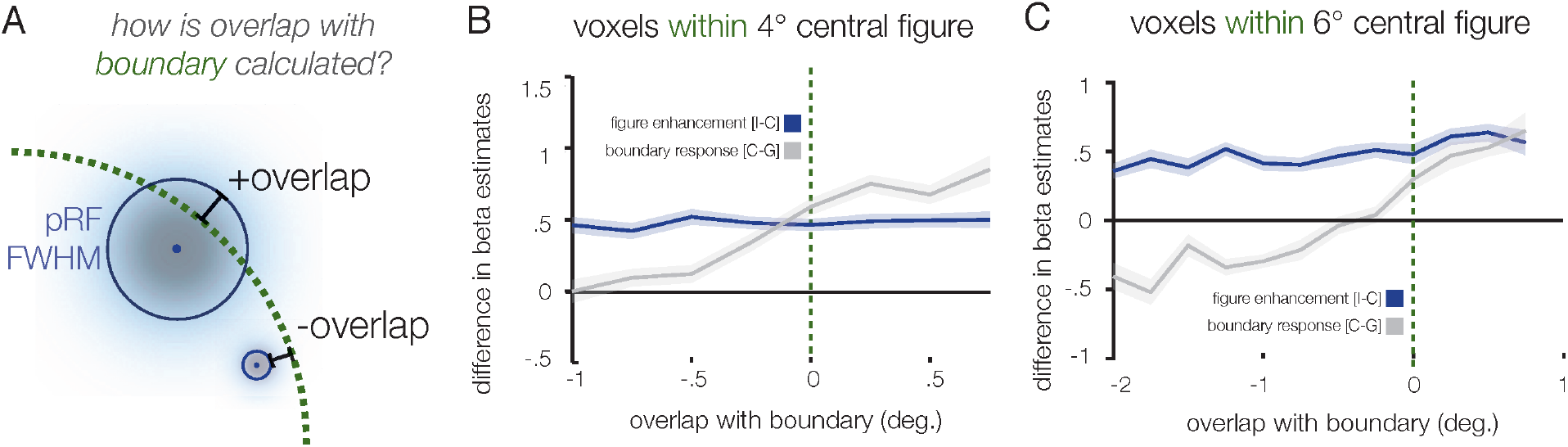
Boundary response and figure enhancement in voxels centered within the figure, plotted as a function of pRF extension beyond boundary. As illustrated in (A), two pRFs with centers at the same eccentricity within the figure might, by virtue of differences in their side, lead to differential sensitivity to the location of the figure-surround boundary. We calculated whether the central FWHM-region of each pRF fell short of the boundary (negative ‘overlap’ values) or extended past the boundary (positive ‘overlap’ values). (B) V1 boundary responses and figure enhancement as a function of distance; while boundary responses (congruent minus ground-only, grey) is evident primarily in voxels whose pRFs overlap with the figure/surround boundary (overlap > 0), figure enhancement (incongruent minus congruent, blue) occurs along the full extent of measured distances. Bins are 0.25° wide, and those containing fewer than 10 data points were trimmed. (C) Results from a control experiment using 6°-diameter figures. While boundary responses again primarily occur in voxels whose pRFs overlap with the boundary (overlap < 0), figure enhancement persists even in voxels with up to 2° spatial separation with the figure-surround boundary.

We observed this same pattern of results in a separate control experiment using larger 6°-diameter figures. Participants again performed a visual monitoring task on the central fixation point (0.5° diameter), so both figure and ground stimuli were task-irrelevant. The greater response to incongruent than congruent figures was evident even in voxels whose pRF response fields (FWHM) were more than 2° away from the figure-surround boundary (Figure 5C; F(1,6) = 1.66, p = .25). In contrast, our measure of the boundary response (congruent minus ground-only) was again local to the border, and showed no evidence of spreading toward the center of the figure. Together, these results imply that the enhancement stemming from the figure’s incongruent orientation does not arise from strictly local processing of feature contrast, as such effects would be expected to decline as a function of distance from the boundary. Instead, the heightened response we observe for incongruent figures, as compared to congruent figures, is consistent with the predicted effects of figure enhancement in V1.

Finally, we used pRF measurements to identify voxels within V1 whose FWHM envelopes were either fully contained within the figure (Figure 6A, orange), overlapped with the boundary (magenta) or fell in the surround region beyond the boundary (purple). Mean BOLD time courses and Beta estimates of fMRI response amplitude revealed a distinct pattern of results for these regions of interest (ROIs). The figure ROI showed a significantly greater response to incongruent figures than to congruent figures (t(5) = 7.86, *p* = 5.4 x 10^-4^), consistent with the predicted effects of figure enhancement, whereas it did not show a boundary effect (t(5) = 0.88, *p* = 0.42).

**Figure 6.**
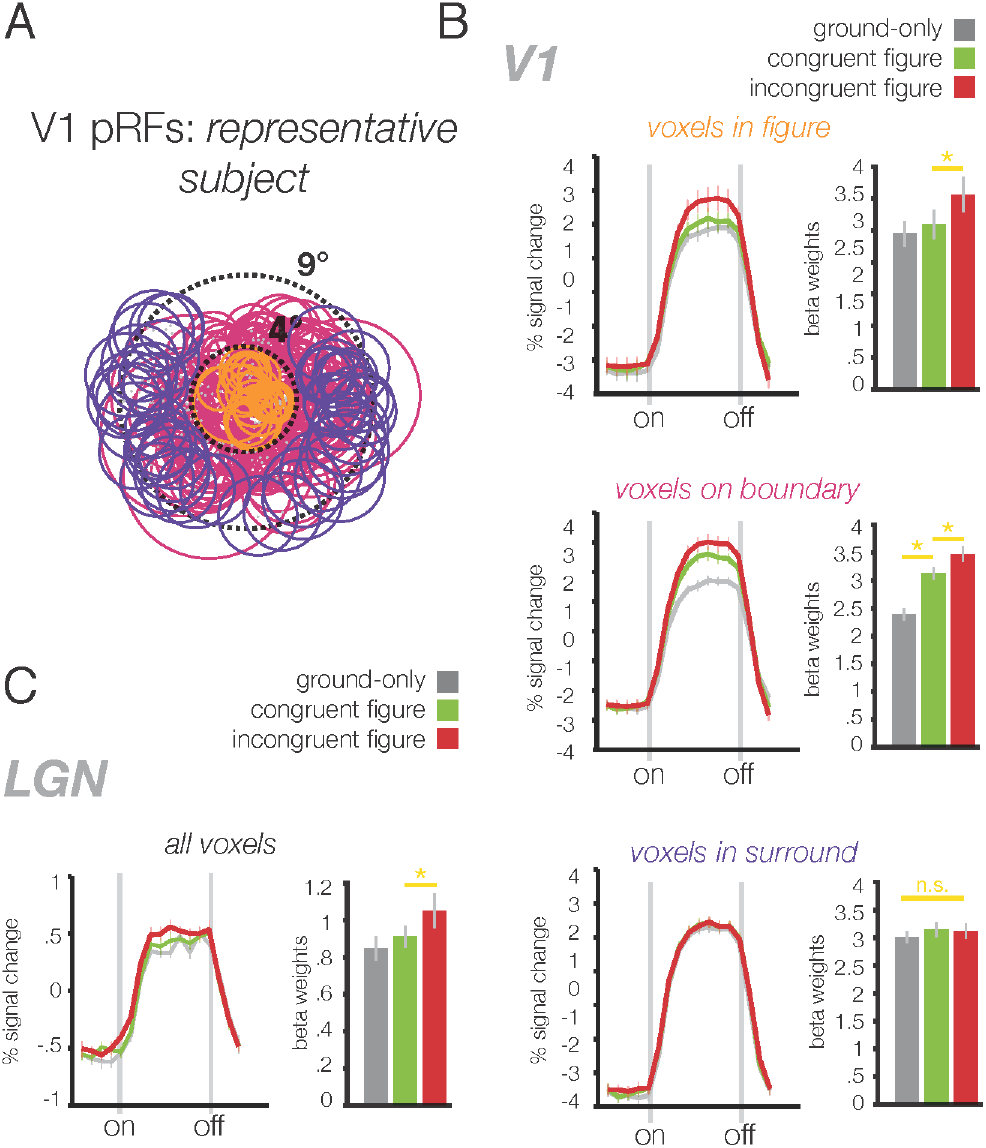
Average BOLD responses across figure, boundary, and surround-selective voxels. (A) V1 pRF locations of one representative subject (circles denote FWHM), illustrating how pRFs were sorted to create ROIs with voxels primarily responsive to the figure (orange, 205 voxels across participants), surround (purple, 293 voxels), or on the boundary (magenta, 476 voxels). Dotted lines denote the spatial extent of our mapping stimulus (9° diameter) and the figure region (4° diameter). (B) V1 results for these ROIs; left panels depict average BOLD time courses in each condition, with dotted lines marking the onset and offset of 16s stimulus blocks. Right panels depict averaged estimated GLM beta weights in each condition. Error bars indicate ±1 SEM between subjects. (C) Results from voxels in the LGN that overlapped with the center, surround, and boundary.

The boundary ROI exhibited a much stronger response to congruent figures than to ground only (t(5) = 7.7, *p* = 7.4 x 10^-4^), consistent with our prediction that a border response would be evoked by the phase-misaligned iso-oriented figure. The boundary ROI also showed a significantly greater response to incongruent than congruent figures (t(5) = 5.46, *p* = 2.8 x 10^-3^), consistent with our expectation that boundary ROI voxels should show some degree of figure enhancement, given that part of their population receptive fields fall upon the figure region. A direct comparison indicated that the magnitude of the boundary response was significantly greater than the magnitude of figure enhancement in these boundary-ROI voxels (t(5) = 3.29, p = .011; mean Beta difference of 0.74 (SE = 0.10) and 0.35 (SE = 0.06), respectively), consistent with findings reported in studies of non-human primates (Poort et al., 2012; Self et al., 2019). Finally, the surround ROI did not show any significant differences between the experimental conditions, further demonstrating the spatial specificity of figure-ground modulation. Taken together, these results provide strong support for the notion that the mechanisms of figure enhancement and sensitivity to the figure-surround boundary lead to distinct patterns of activation in human V1.

### Reconstructing differential responses to figures in stimulus space

pRF modeling also provides an opportunity to visualize the pattern of voxel responses for a given visual area in the native stimulus space (cf. Kok and de Lange, 2014). Here, we used regularized linear regression to infer the spatial profiles of both boundary responses and figure enhancement responses in stimulus space. As described in Figure 7, this approach assumes that a voxel-wise vector of response amplitudes can be modeled as a linear product of those voxels’ pRFs and the spatial profile of the differential response in stimulus space. To improve the model’s tractability, we downsampled the effective resolution of the stimulus space to 4 pixels per degree of visual angle, such that each pixel in the reconstructed spatial profile (Figure 7) corresponds to 0.25° of the original display.

**Figure 7.**
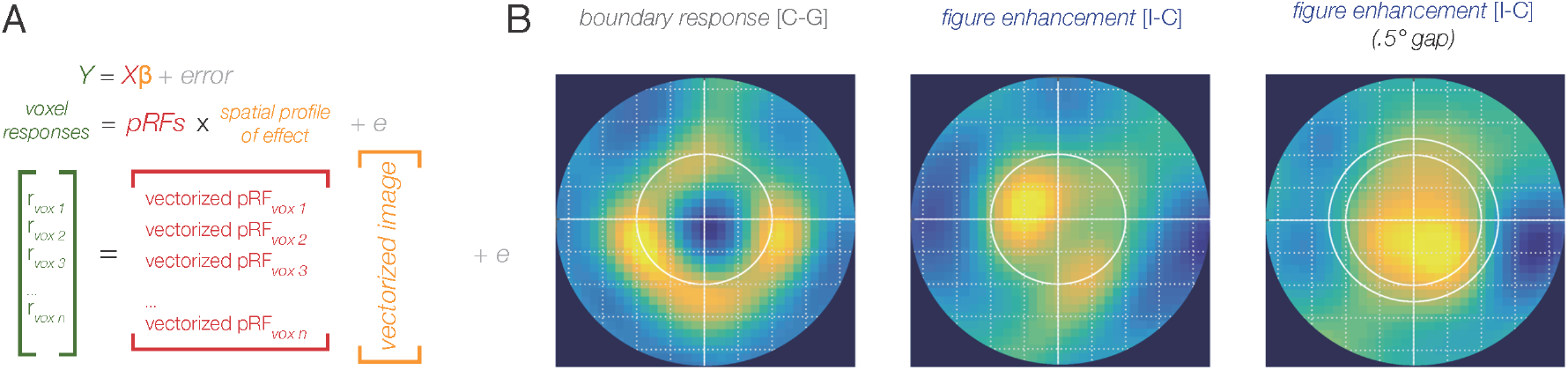
pRF-based visualization of voxelwise BOLD responses in stimulus space. (A) Schematic of the multivariate regression-based method of estimation; the resulting images are reshaped from the estimated predictor term (yellow). The precise formula used appears in the Methods. (B) Panels indicate the projected spatial profiles of the differential responses associated with boundary response (congruent minus ground-only, left) and figure enhancement (incongruent minus congruent) for V1 data in the main experiment (middle) and in a control experiment which introduced a 0.5° gap between figure and surround (right). Each pixel corresponds to 0.25° of visual angle of the original stimulus space; contrast depicts predictor values, normalized and averaged across subject. See also Figure 9.

We used an L2 (ridge) penalty term to minimize overfitting (Hoerl and Kennard, 1970), and ten-fold cross-validation to select the penalty value that produced a minimal RMSE. This regression method can yield better spatial resolution in the visualization than simply scaling the 2D Gaussian derived from each voxel’s pRF by its response magnitude and averaging the weighted responses of all voxel pRFs throughout the visual field, as the latter approach is sure to introduce some blurring of the reconstruction given the spatial spread of the Gaussian pRFs themselves. We performed this analysis on the differential response to congruent figures versus ground-only and to incongruent versus congruent figures, normalized the resulting images within subjects, and then created averaged visualizations across subjects as shown in Figure 7. The spatial profile of these visualization results clearly support the distinct effects of boundary responses and figure enhancement, and go on to show that the response pattern evoked by incongruent figures, as compared to congruent figures, is well described by enhanced representation specific to the figural region.

### A visible local boundary is not sufficient for figure enhancement in V1

We have demonstrated that orientation-defined figures produce enhanced BOLD responses across the figure region in V1, and that this enhancement is spatially distinct from responses associated with local processing of a phase-defined boundary. We ran an additional control experiment to ascertain whether the presence of a clearly visible boundary, rather than the orientation difference between figure and surround, might be sufficient to induce an effect of figure enhancement in V1. This control relied on the same stimulus configuration and task but introduced a 0.5° gap (2.0-2.5° eccentricity) between the figure and the surround (Figure 8), so that the boundary between figure and surround could be very clearly perceived in both congruent and incongruent conditions. The reconstruction of V1 activity again shows that the orientation difference between figure and surround led to enhanced responses throughout the extent of the central figure (Figure 7B, right). Thus, we can conclude that figure enhancement cannot be explained simply in terms of an inward spread of local boundary responses; larger-scale differences in feature content that distinguish the figure from the surround confer an additional enhancement.

**Figure 8.**
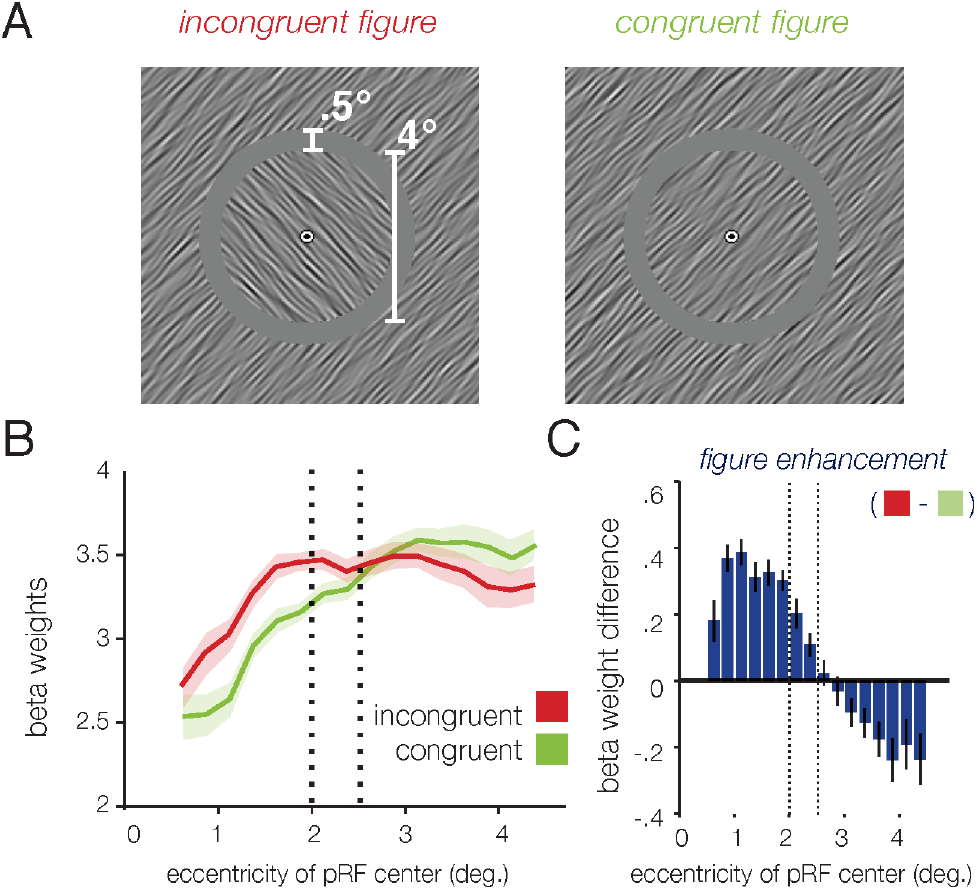
Stimuli (A) and results (B,C) of Control Experiment A, in which a 0.5° greyscale gap separated the figure and surround. Plotting follows conventions of Figure 2; here, since both conditions have a visible boundary, we can subtract responses of the congruent figure condition (green) from the incongruent figure condition (red) to yield orientation-dependent figure enhancement. Results closely follow the pattern of the main experiment, with the observed enhancement extending throughout the figure region.

### Results in areas V2 and V3

Our pRF mapping experiments indicated that pRF sizes were somewhat larger in V2 and V3 (see Figure 4A), as has been previously reported (Dumoulin and Wandell, 2008; Wandell and Winawer, 2015). Nevertheless, the pattern of results in V2 and V3 were very similar to those found in V1, in accord with recent intracranial recordings of human V2/V3 neurons in response to feature-defined figures (Self et al., 2016). Note that the larger peripheral pRFs found in these extrastriate visual areas are more likely to encroach on the 2° figure-surround boundary, and thus border response effects are evident in a larger proportion of voxels than in V1. We again averaged the responses across voxels whose pRFs fell primarily in the figure, in the surround, or overlapped the boundary, assuming a FWHM central region. Voxels with pRFs confined well within the figure showed significant effects of figure enhancement (V2: t(5) = 5.37, *p* = 0.0030; V3: t(5) = 4.81, *p* = 0.0048) but did not show a significant boundary response (V2: t(5) = 1.38, *p* = 0.23; V3: t(5) = 1.12, *p* = 0.31). Voxels whose pRFs fell on the boundary exhibited both effects (V2: t’s(5) > 5.16, *p’s* < 0.0036; V3: t’s(5) > 4.27, *p’s* < 0.0079). In V2, voxels whose pRF envelopes were contained within the surround did not show evidence of either type of enhancement (V2: t’s(5) < 1.64, *p’s* > 0.16). In V3, voxels corresponding to the surround also showed no effect of figure enhancement (t(5) = 0.46, *p* = 0.80), though they did show a significant effect of boundary response (t(5) = 2.91, *p* = .034). Of note, however, was that pRFs in this peripheral region of V3 were quite large and more spatially diffuse in their response profile; only 96 voxels across six participants were included in the surround-only subset of voxels.

### Figure-ground modulation in the LGN

While our primary aim was to resolve the spatial extent of figure-ground modulation in the early visual cortex, we were also able to obtain measures of BOLD activity in the lateral geniculate nucleus (LGN). In recent work, we have shown that the LGN is sensitive to figure-ground modulation in the absence of directed attention (Poltoratski et al., 2019). Although the receptive fields of individual LGN neurons are small, each fMRI voxel covers a proportionally greater segment of this small subcortical structure than a similar voxel sampled from cortex. As such, the pRFs that we measured in the LGN were too large to clearly distinguish responses to the central figure region from those to the surround (eccentricity 2-4.5°); 96.2% of recorded voxels overlapped with the figure-surround boundary. However, we did find evidence of figure enhancement (incongruent > congruent figure, t(5) = 2.89, *p* = 0.034) across the full set of LGN voxels as shown in Figure 6C; there was no significant difference between responses in the congruent figure condition versus ground-only (t(5) = 2.27, *p* = 0.072).

We also noted a similar pattern of results in Control Experiment B, in which a larger figure was shown. When we analyzed the response of 50 voxels in this experiment whose pRFs fell on the figure region but also extended beyond the boundary, we observed significantly greater responses to incongruent figures than to congruent figures, (t(2) = 8.2, *p* = 0.014), consistent with our predicted effects of figure enhancement. In contrast, this LGN ROI did not show a significant difference in its response to congruent figures when compared to ground-only (t(2) = 1.32, *p* = 0.32), indicating a lack of sensitivity to the phase-defined border present in the congruent figure condition. We then performed a more spatially restricted analysis, focusing on the 8 voxels whose pRFs fell within the figure region, and again found a significant effect of figure enhancement (t(2) = 4.7, *p* = .04). These results suggest that activity in the lateral geniculate nucleus is involved in the broader visual circuitry that supports figure-ground perception, presumably via top-down feedback from the visual cortex. The current results serve to replicate and extend our previous findings (Poltoratski et al., 2019) and other recent work (Jones et al., 2015) describing the modulation of LGN responses by figure-ground configurations.

## Discussion

Our study provides compelling evidence that early visual areas in the human brain are engaged in figure-ground processing; they respond to local boundary differences, but confer an additional enhancement of the entire figure region for figures that differ in featural content from their surround. We demonstrate that that this figure enhancement is spatially distinct from local responses associated with sensitivity to the boundary using a novel response reconstruction method informed by pRF mapping of voxels in the early visual cortex. While a phase-defined border was present in the congruent figure condition, this did not lead to widespread effects of figure enhancement, consistent with the weaker perceptual salience of the figure. It was also the case that figure enhancement occurred when the local orientation-defined boundary was obscured by introducing a 0.5° gap between the figure and surround. Our results suggest that the presence of a local feature-defined boundary is neither necessary nor sufficient to observe V1 enhancement to figures with distinct orientation content. Boundary enhancement is predicted by several known mechanisms, including a release from feature-tuned surround suppression that serves to enhance local orientation contrast (Blakemore and Tobin, 1972; Nelson and Frost, 1978; Nothdurft, 1991; Sillito et al., 1995; Bair et al., 2003; Shushruth et al., 2012) or sensitivity to high spatial frequency information at the boundary (Landy and Kojima, 2001; Mazer et al., 2002; Hallum et al., 2011). However, the asymmetric enhancement of the central figure region is not readily attributable to these well-known neural mechanisms associated with local processing within V1.

Unlike boundary detection, which serves to enhance local feature differences and presumably involves local processing within V1 (Self et al., 2013; Bijanzadeh et al., 2018), figure enhancement appears to serve a more integrative function of grouping regions that *share* one or more features that distinguish it from the surround. Compelling neurophysiological work in the monkey has pointed to the role of feedback from higher cortical areas, including V4 (Self et al., 2012; Klink et al., 2017), as critical for observing effects of figure enhancement but not boundary detection in V1 (Lamme and Roelfsema, 2000). Enhanced responses to the border between figure and surround emerge rapidly in V1 neurons, and are evident in the initial transient response, whereas figure enhancement emerges later in time and remains sustained over longer viewing periods (Lamme and Roelfsema, 2000; Scholte et al., 2008; Poort et al., 2012). Accordingly, boundary detection and figure enhancement appear to modulate different laminar layers of V1, with the latter primarily leading to enhanced responses in layers 1,2, and 5, which receive feedback from higher visual areas (Self et al., 2013). While fMRI does not provide information about the source of the modulations we observe here, our results are generally supportive of these proposed mechanisms of feedback. For voxels with pRFs that overlapped the figure, we found no relationship between the proximity of a voxel’s response field to the boundary and the magnitude of figure enhancement when contrasting responses to incongruent and congruent figures. Our findings deviate from standard accounts of orientation-tuned surround suppression, in which local inhibitory interactions lead to suppression that falls off symmetrically as a function of spatial separation between a target stimulus and the surround stimulus (Bair et al., 2003; Bijanzadeh et al., 2018). They also run counter to alternative theories that challenge the need for mechanisms of figure enhancement beyond detection of the boundary (c.f. Rossi et al., 2001; Zhaoping, 2003). Previous studies were not able to characterize the spatial profile of these distinct mechanisms in the human as we were able to carry out here using high-field imaging, pRF modeling, and computational methods for regression-based visualization.

To date, figure processing in humans has been largely studied in higher visual areas, including V4 (Kastner et al., 2000; Kourtzi et al., 2003; Thielscher et al., 2008) and the lateral occipital cortex (Grill-Spector et al., 1999; Vinberg and Grill-Spector, 2008); indeed, some early neuroimaging work did not find modulations in V1 in response to feature-defined textures or figures, likely due to signal strength or sensitivity (Kastner et al., 2000; Schira et al., 2004). However, this work and others (Appelbaum et al., 2008; Scholte et al., 2008; Self et al., 2016; Poltoratski et al., 2019) provide convergent evidence of common mechanisms of figure-ground processing across human and non-human primates, in which processing in higher-level visual areas including V4 informs, via feedback, enhancement of responses to figure regions in V1.

This work builds upon several recent findings of V1 contributions to mechanisms of visual segmentation and grouping, including visual salience (Zhang et al., 2012; Poltoratski et al., 2017), grouping (Murray et al., 2002; Kourtzi et al., 2003; Beck and Kastner, 2005; Roelfsema, 2006), perceptual filling-in (Sasaki and Watanabe, 2004; Meng et al., 2005; Hong and Tong, 2017), and the processing of illusory surfaces (Kok and de Lange, 2014; Kuai et al., 2017). Together, these studies point to an important role of the early visual system in figure perception through a combination of both local feedforward mechanisms and automatic feedback mechanisms, which provide contextually-driven modulation of responses in the absence of directed attention or particular task demands. While it may be the case that attention differentially modulates figure enhancement and boundary detection (Lamme et al., 1998b; Qiu et al., 2007; Poort et al., 2012; Self et al., 2019), consistent with their separable mechanistic origins, growing evidence suggests that figure enhancement is not a simple byproduct of spatial attention (Marcus and van Essen, 2002; Jones et al., 2015; Papale et al., 2018; Poltoratski et al., 2019). Indeed, it appears that both humans and monkeys exhibit spatially specific effects of figure enhancement irrespective of the locus of attention, task, or training exposure to feature-defined figures (Poltoratski et al., 2019; Self et al., 2019).

This study also highlights a useful and intuitive method for utilizing population receptive field modeling to reconstruct visual responses (Thirion et al., 2006; Miyawaki et al., 2008; Naselaris et al., 2009; Kok and de Lange, 2014), which can move toward bridging the gap between the resolution of spatial effects measured by neurophysiological recordings and human neuroimaging. The regression method allows for a finer spatial resolution of reconstruction than a simple summation of pRF responses weighted by their response amplitudes (Figure 9). The latter approach is necessarily constrained by the resolution of pRFs themselves: even perfectly noise-free data will yield blurry reconstructions if the Gaussian pRFs are large, relative to the stimulus, and greater effects of pRF blurring will occur when ascending the visual hierarchy. This regression-based reconstruction approach could be adopted more widely to estimate the spatial profile of fMRI BOLD effects at spatial scales that would prove challenging with standard approaches.

**Figure 9.**
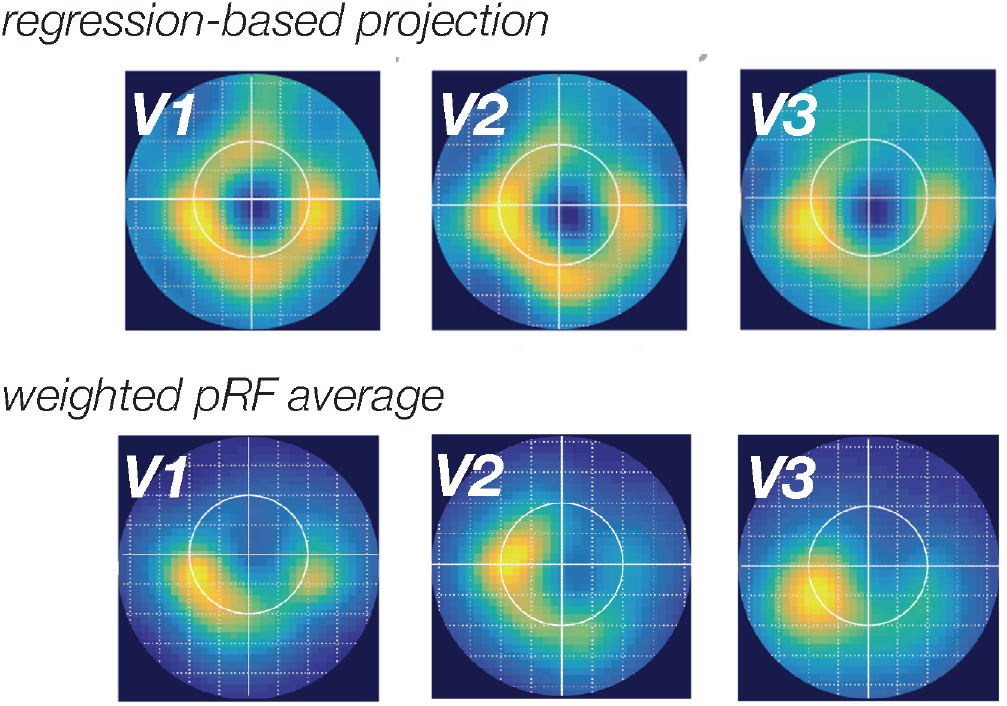
A comparison of two methods for visualization of the boundary response (congruent minus baseline) in areas V1, V2 and V3. The left panel shows our ridge regression-based approach, while the right panel is a simpler average of each pRF weighted by the BOLD amplitude of boundary response associated BOLD Beta weight difference in that voxel. The regression-based approach generally led to better resolved reconstructions, particularly in higher visual areas which have typically larger pRFs.

## Acknowledgements

This work was supported by the National Institutes of Health (Grant R01EY029278 to FT), National Science Foundation (Grant BSC-1228526 to FT), a National Science Foundation Graduate Research Fellowship to SP; and the National Institutes of Health (Grant P30-EY008126) center grant to the Vanderbilt Vision Research Center.

---

<u>Movie 1.</u> *Illustration of the three main experimental stimuli conditions: incongruent figure, congruent figure, and ground-only*. Timing is truncated for the illustration: in the experiment, stimulus blocks were 16s long, and were interspersed with 16s fixation-rest periods. Orientations (45° and 135°) were counterbalanced across blocks, and timing was randomized for every experimental run and participant. Participants reported the brief 200ms color change at fixation throughout the entire run.

<u>Movie 2.</u> *Illustration of the experimental conditions of Control Experiment A, in which a 0.5° greyscale gap was introduced*. As in Movie 1, timing is truncated for this illustration.

## Notes

The authors declare no competing financial interests.

